# Platform for rapid nanobody discovery *in vitro*

**DOI:** 10.1101/151043

**Authors:** Conor McMahon, Alexander S. Baier, Sanduo Zheng, Roberta Pascolutti, Janice X. Ong, Sarah C. Erlandson, Daniel Hilger, Aaron M. Ring, Aashish Manglik, Andrew C. Kruse

## Abstract

Camelid single-domain antibody fragments (“nanobodies”) provide the remarkable specificity of antibodies within a single immunoglobulin V_HH_ domain. This unique feature enables applications ranging from their use as biochemical tools to therapeutic agents. Virtually all nanobodies reported to date have been obtained by animal immunization, a bottleneck restricting many applications of this technology. To solve this problem, we developed a fully *in vitro* platform for nanobody discovery based on yeast surface display of a synthetic nanobody scaffold. This platform provides a facile and cost-effective method for rapidly isolating nanobodies targeting a diverse range of antigens. We provide a blueprint for identifying nanobodies starting from both purified and non-purified antigens, and in addition, we demonstrate application of the platform to discover rare conformationally-selective nanobodies to a lipid flippase and a G protein-coupled receptor. To facilitate broad deployment of this platform, we have made the library and all associated protocols publicly available.

## Introduction

Antibodies have had a transformative impact on science and medicine due to their exceptional specificity and biochemical versatility, enabling applications in almost every aspect of biomedical inquiry. Conventional antibodies are composed of two heavy chains and two light chains. Each chain contributes to antigen binding specificity through a variable domain, termed V_H_ and V_L_ for the heavy and light chain, respectively. A key exception to this general architecture is found in camelids (llamas, alpacas, and their relatives), which possess a parallel antibody repertoire composed solely of heavy chains^1,2^. Such antibodies bind to their target antigens through a single variable domain, termed V_HH_, which contains the entire antigen-binding surface. Unlike the antigen binding fragments of conventional antibodies (Fabs), isolated V_HH_ domains (also called “nanobodies”) can be readily expressed in bacteria as the product of a single gene, and in many cases these fragments can even fold and retain antigen specificity in the reducing environment of the cytosol. Owing to their versatility, nanobodies have found applications in protein biochemistry and structural biology, cell biology, and as potential diagnostic and therapeutic agents^2–6^.

Despite the growing importance of nanobodies throughout biomedical research, the current methods for creating nanobodies remain slow, costly, and often unreliable. The majority of nanobodies described to date have been derived from immunization of camelids, a lengthy and expensive process posing a significant barrier to entry for most laboratories. Furthermore, animal-derived antibodies are frequently restricted from binding conserved epitopes due to immunological tolerance of self-antigens. Other efforts have sought to address these challenges by combining phage display with a synthetic nanobody library^7^. However, these libraries remain restricted to contract work with a commercial provider, limiting their broad utilization. More importantly, with synthetic libraries it remains particularly challenging to identify nanobodies that not only bind to their target, but also specifically recognize a defined conformation – this represents one of the most important applications of animal derived nanobodies. Phage display techniques have proven useful in isolating binders, but functional characterization is largely dependent on recombinant expression and purification. Here, we address these challenges through the development and application of a fully synthetic nanobody library for rapid discovery of novel affinity reagents using yeast surface display. Importantly, the use of a cell-based selection scheme allows straightforward identification of conformationally-selective nanobodies for challenging membrane protein targets and even non-purified protein samples. To enable broad utility of this new resource, we have made the library and all associated protocols available to the scientific community.

## Library construction and validation

To develop a platform for rapid *in vitro* nanobody discovery, we first sought to design a synthetic nanobody library starting from a consensus framework derived from llama genes IGHV1S1-S5. This constant framework was combined with designed variation of the complementarity determining loops (CDRs) that comprise the highly variable, antigen-binding interface of the V_HH_. Although there are many previously described strategies for introducing variation in antibody CDR loops ^8^, we postulated that nanobodies of known structure in the Protein Data Bank (PDB) represented an especially curated set of highly stable, biochemically well behaved variants as evidenced by their tractability in crystallographic studies. The entire set of unique nanobodies in the PDB (93 sequences at the time of analysis) was analyzed for position specific variation in the CDRs, and our design aimed to recapitulate this diversity. For the residues immediately adjacent to the CDRs, we introduced partial randomization, allowing only a few possible amino acids guided by the observed frequencies of nanobodies in the PDB. For the highly variable positions in each CDR, we introduced much more aggressive randomization. First, the frequency of each amino acid within CDR3 was determined for all nanobodies in the PDB. The resulting mixture of amino acids was modified to eliminate cysteine and methionine to avoid chemical reactivity, and was introduced into various positions in each CDR in the synthetic library. Based on analysis of nanobody sequences, we elected to introduce four such highly variable positions in CDR1 and one in CDR2. The longer CDR3 represents a critically important loop for antigen recognition – here we introduced either seven, eleven, or fifteen consecutive positions of high diversity mixture to emulate CDR3 length variation seen in the natural repertoire (Figure 1A – C).

**Figure 1.**
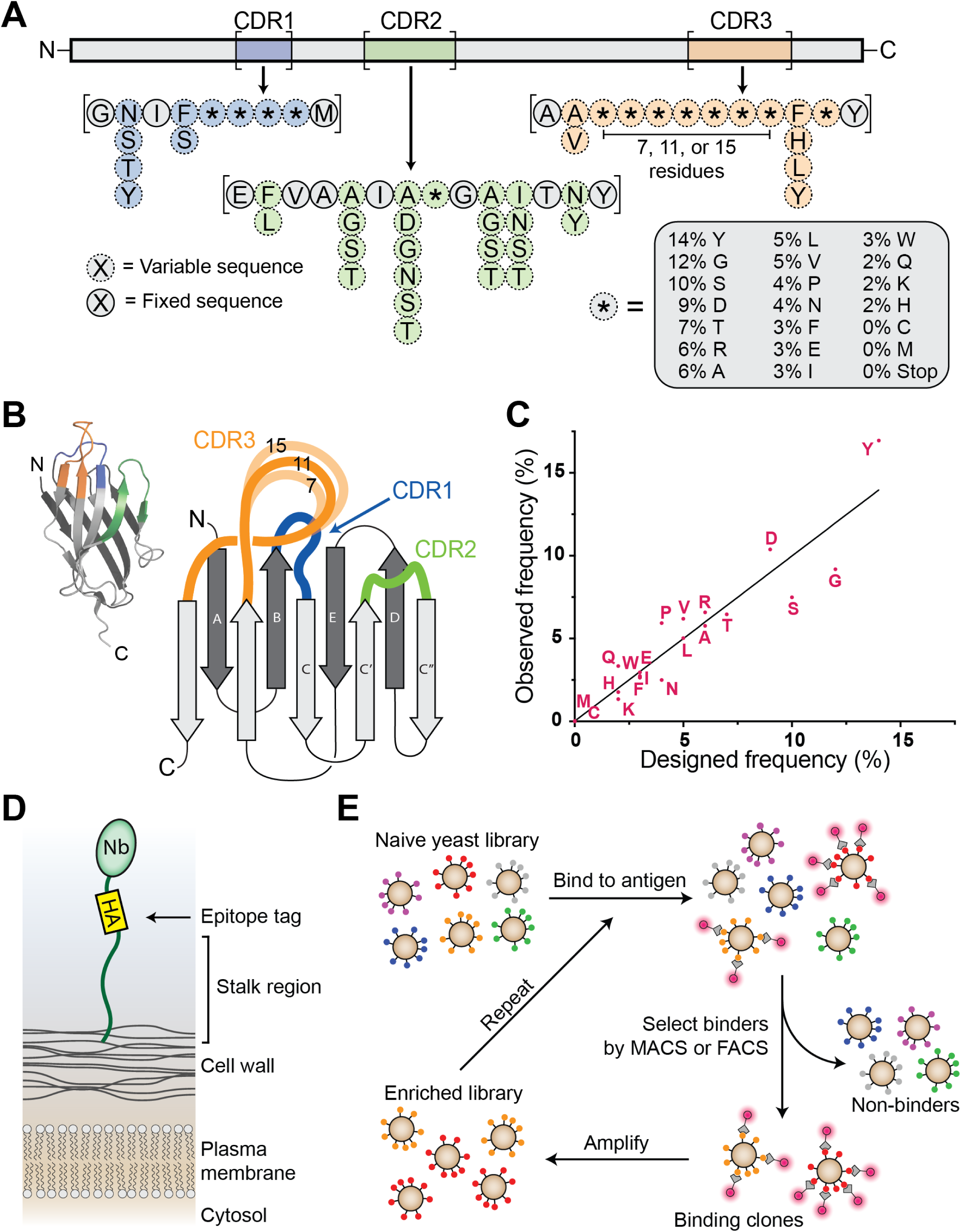
Design and construction of synthetic nanobody library. (A) Schematic of synthetic nanobody. Framework regions were fixed in sequence (gray), while portions of the CDR loops were varied (blue, green, and orange). Partial randomization was achieved with mixed nucleotides to allow up to six possible residues (dotted lines), while highly variable regions were synthesized using a trimer phosphoramidite mixture (asterisks with dotted outlines). (B) Overview of a nanobody showing three-dimensional structure (inset) as well as a cartoon schematic highlighting CDR loops. Synthetic CDR3 sequences used variable lengths of 7, 11, or 15 residues of trimer phosphoramidite mixture. (C) Amino acid frequencies of diversified positions in CDRs were analyzed by next-generation sequencing (NGS), showing that library frequencies closely matched target values. (D) Schematic of nanobody display on yeast. Nanobody is shown with an HA tag at the C-terminus followed by a long flexible stalk tethering the nanobody to the yeast cell wall. (E) Schematic of the nanobody selection process. Antigen is shown with a fluorescent tag as a glowing red circle. Yeast displaying nanobodies with affinity to antigen are shown being isolated via MACS or FACS, amplified, and undergoing iterative rounds of selection. See also Figures S1 and S2, Table S1.

Primers encompassing the desired sequence diversity were synthesized, with high diversity regions constructed using a mixed pool of trimer phosphoramidites to match the target codon frequency ^9,10^. We then built a pooled DNA library encoding the full nanobody sequence by assembly PCR of these primers (see Methods for full details). A pervasive but often underappreciated drawback of synthetic antibody fragments is poor biochemical behavior. Prior to generating a large yeast display library, we sought to assess whether our library design would reliably produce biochemically tractable clones suitable for structural and cell biological studies. A total of 11 nanobody sequences were chosen at random from the library DNA and tested for biochemical behavior by expression and purification from *Escherichia coli*. Of these clones, 9 were well expressed (>20 mg/L), could be purified by routine procedures, and showed reasonably monodisperse size exclusion profiles (**Figure S1**; **Table S1**).

Next, we sought to develop a straightforward way to tether nanobodies and other small proteins to the surface of *Saccharomyces cerevisiae* and other yeast species. A variety of methods for this purpose have been reported previously, with the most common approach being fusion of a protein of interest to Aga2p, a yeast cell wall protein^11^. While this strategy has been widely used, it is limited by the requirement to use specialized yeast strains expressing galactose-inducible Aga1p, which serves to tether Aga2p to the yeast cell wall. Instead, we built a simplified system in which the protein of interest is directly connected to the cell wall through a single tether designed to replace the Aga2p-Aga1p linker protein. A synthetic tether mimicking the low complexity sequence of yeast cell wall proteins was designed, and was evaluated for its ability to anchor a test protein to the yeast cell wall (see **Supplementary Protocol** for full sequence). Remarkably, accessibility of this protein correlated strongly with the molecular weight of the staining reagent, suggesting steric occlusion by cell wall glycans. Consistent with this explanation, longer tethers alleviated this problem, and above 600 amino acids in length, molecular weight dependence of staining levels was negligible (**Figure S2**). For library creation, we chose to use a 649 amino acid tether. In addition, we included an N-terminal engineered mating factor α pre-protein which has been reported previously to enhance antibody expression in yeast^12^. On the C-terminus, we included a glycosylphosphatidylinositol anchor sequence, which results in covalent tethering of the protein to the yeast cell wall^13^ (Figure 1D). On the basis of these results, the engineered surface display plasmid (pYDS649) was linearized and then transformed into *Saccharomyces cerevisiae* protease deficient strain BJ5465 together with the DNA library, which had been amplified to include flanking sequences homologous to pYDS649 for recombination. This resulted in a yield of 5 × 10^8^ transformants.

To characterize the resulting yeast surface displayed library, we isolated plasmid DNA from ~600,000 yeast cells and performed high throughput sequencing. Analysis of 480,000 unique sequences confirmed that the library clones possessed the desired amino acid frequencies in the hypervariable portions of the CDR loops (Figure 1C), and also confirmed that CDR-flanking region diversity resembled target frequencies. Overall, 24% of sequences analyzed were full length and free of frameshift errors introduced during the assembly PCR process. This value aligned closely with the observation that 26% of library yeast showed high expression of nanobodies as measured by flow cytometry. Taken together, these data imply that the library contains a total of approximately 10^8^ unique full-length nanobody clones that are expressed and displayed on the yeast surface.

## Identification of nanobodies using magnetic cell sorting

We next sought to determine if our approach could yield nanobody clones targeting diverse antigen classes, including both soluble and membrane proteins. As an initial test, we aimed to identify nanobodies targeting human serum albumin (HSA), the most abundant protein in human plasma. Binding of therapeutic proteins and small molecules to HSA is a key determinant of drug pharmacokinetics; nanobodies targeting HSA may therefore provide a modular tool for enhancing the half-life of biologics^14–16^. In this first test, we exclusively employed magnetic cell sorting (MACS), a low-cost cell separation method that can be performed in basic laboratory environments without specialized equipment^17^.

To identify novel HSA-binding nanobodies, we first prepared fluorescently-labeled HSA protein and then used this reagent to perform iterative staining and magnetic-bead based enrichments. In brief, this approach entails incubation of the yeast library with fluorescent HSA antigen, washing off excess antigen, and further staining desired clones with anti-fluorophore magnetic microbeads. Labeled cells are then isolated by magnet-based separation and amplified in standard yeast culture medium (Figure 1E). As with any library-based selection, it is of paramount importance to ensure binders recognize the antigen itself, rather than the reagents or fluorescent tags used for separation. To achieve such specificity, yeast with reactivity to beads alone were depleted from the library prior to each selection step. To decrease the probability of enriching fluorophore binders, the fluorophore tag used was alternated between Alexa Fluor 647 and fluorescein isothiocyanate (FITC) for each consecutive round of magnetic selection. After four rounds of selection (Figure 2A), single yeast colonies were isolated and stained with fluorescently labeled HSA for validation by analytical flow cytometry. For further study, we selected one clone with strong antigen binding (named Nb.b201) for in-depth characterization.

**Figure 2.**
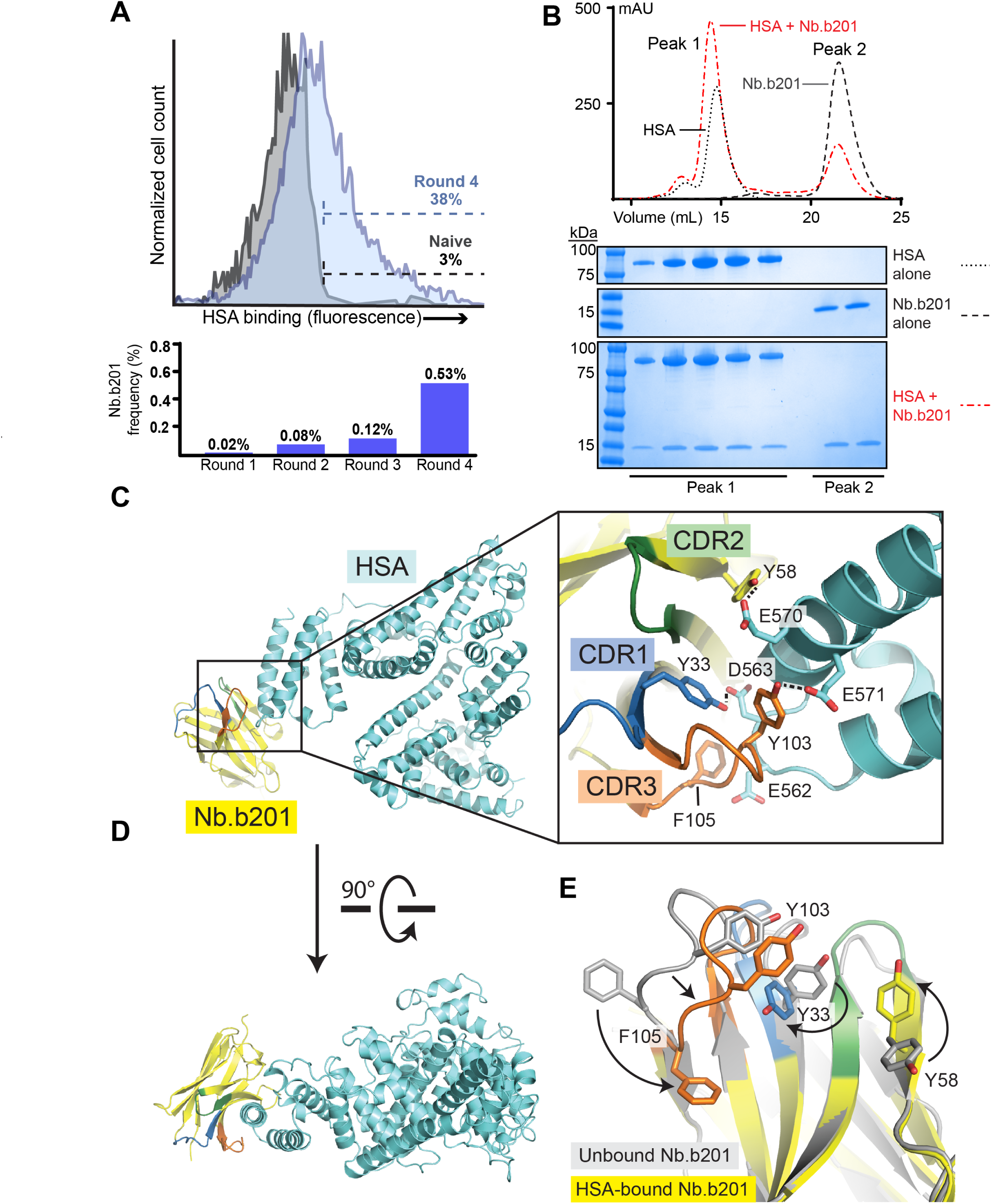
Validation of nanobody platform using human serum albumin (HSA). (A) Efficiency of nanobody selection was assessed by flow cytometry and by NGS, revealing gradual enrichment of HSA binding clones over the course of four selection cycles. (B) Size exclusion chromatography analysis with purified Nb.b201 confirmed binding to antigen. (C) Crystal structure of nanobody:HSA complex PDB ID: 5VNW, including closeup view of CDR loops interacting with antigen. Hydrogen bonds are shown as dotted lines. (D) Rotated PDB ID: 5VNV view of the complex (E) Comparison of nanobody structures in the antigen-bound (yellow) and antigen-free state (gray), showing conformational change upon antigen binding. Changes are highlighted with arrows, and CDR loops are colored blue, green, and orange for CDRs 1, 2, and 3, respectively. See also Figure S3, Table S2.

As expected, Nb.b201 could be expressed and purified from *E. coli* using standard methods, with a yield of 26 mg of purified protein per liter of culture. We assessed HSA binding *in vitro* by analytical size exclusion chromatography (Figure 2B) and more quantitatively by surface plasmon resonance (SPR, **Figure S3**). Both techniques confirmed specific binding of Nb.b201 to HSA. While the binding affinity was modest at 430 nM, Nb.b201 was highly specific, demonstrating no binding to the closely related mouse serum albumin (MSA, **Figure S3**), and forming a stable complex with HSA as assessed by size exclusion chromatography (Figure 2B).

A major goal of our approach presented here is to facilitate structural biology efforts, so we assessed the structural basis for Nb.b201 binding to HSA. Nb.b201 crystallized readily, both in isolation and in complex with HSA. X-ray data collection yielded datasets to a resolution of 1.4 Å and 2.6 Å for the free nanobody and HSA complex, respectively. These crystal structures revealed that Nb.b201 adopts the typical V-set immunoglobulin fold previously observed for animal-derived nanobodies (Figure 2C - E). Comparison of the two structures show provides an opportunity to compare bound and free states of the same antibody fragment. Remarkably, Nb.b201 undergoes large-scale conformational changes upon antigen binding, resulting in the reorientation of the CDR3 backbone and amino acid side chains (Figure 2E). The mode of antigen recognition involves Nb.b201 binding its target largely through its CDR3 loop, with relatively few direct contacts between HSA and CDR1 or CDR2. Interestingly, the epitope recognized by Nb.b201 is a convex protrusion on the surface of HSA, in contrast to most nanobody epitopes, which are more typically concave. The interaction mode includes a mix of polar and non-polar interactions, with a recurring theme of Tyr-Glu and Tyr-Asp hydrogen bonds, an interaction that appears three times in the interface, with the nanobody contributing the Tyr in each case (Figure 2C). These results not only reveal the mechanisms of synthetic nanobody-antigen interaction, but also confirm that useful nanobodies for crystallographic applications can be obtained in a matter of 2 – 3 weeks without use of specialized equipment or large animal immunization.

## Nanobodies targeting non-purified antigen

Selection of yeast or phage surface-displayed libraries is usually performed with purified and labeled protein samples. Protein purification is a highly idiosyncratic process that must be optimized for each individual protein. Even small changes in a variety of factors including pH, ionic strength, presence or absence of cofactors, or detergent concentration can strongly affect protein solubility and activity. As a result, purification and labeling remains a critical bottleneck for protein engineering by surface display methods. This bottleneck poses an even greater barrier for poorly expressed proteins, such as extensively post-translationally modified protein hormones. To assess the versatility of our yeast-based nanobody discovery platform, we next sought to develop methods enabling the use of non-purified antigens as selection reagents.

We focused our efforts on the metabolic hormone adiponectin, a complex protein with many post-translational modifications including O-linked glycosylation and proline hydroxylation, among others (Figure 3). Full-length adiponectin can only be expressed in eukaryotic cells, and even in this case yields are typically low. Human adiponectin tagged with an amino-terminal FLAG epitope was expressed as a secreted protein in HEK293 cells, and the resulting conditioned medium containing a complex mixture of proteins and small molecules was used as the selection antigen (Figure 3A, B). Library-expressing yeast were incubated with adiponectin-containing conditioned medium, washed to remove excess unbound proteins, and then stained with fluorescently-labeled anti-FLAG M1 antibody. Adiponectin-binding yeast were isolated after three rounds of MACS followed by fluorescence activated cell sorting (FACS; Figure 3C). Adiponectin binding was confirmed for three unique clones by SPR, with the best clone (Nb.AQ103) displaying an affinity of 510 nM. These are the first specific nanobodies targeting adiponectin, and broader application of this strategy may enable reagent development for many secreted and transmembrane proteins. Approaches like these may enable rapid reagent development for the large fraction of the human proteome that remains poorly characterized by biochemical methods.

**Figure 3.**
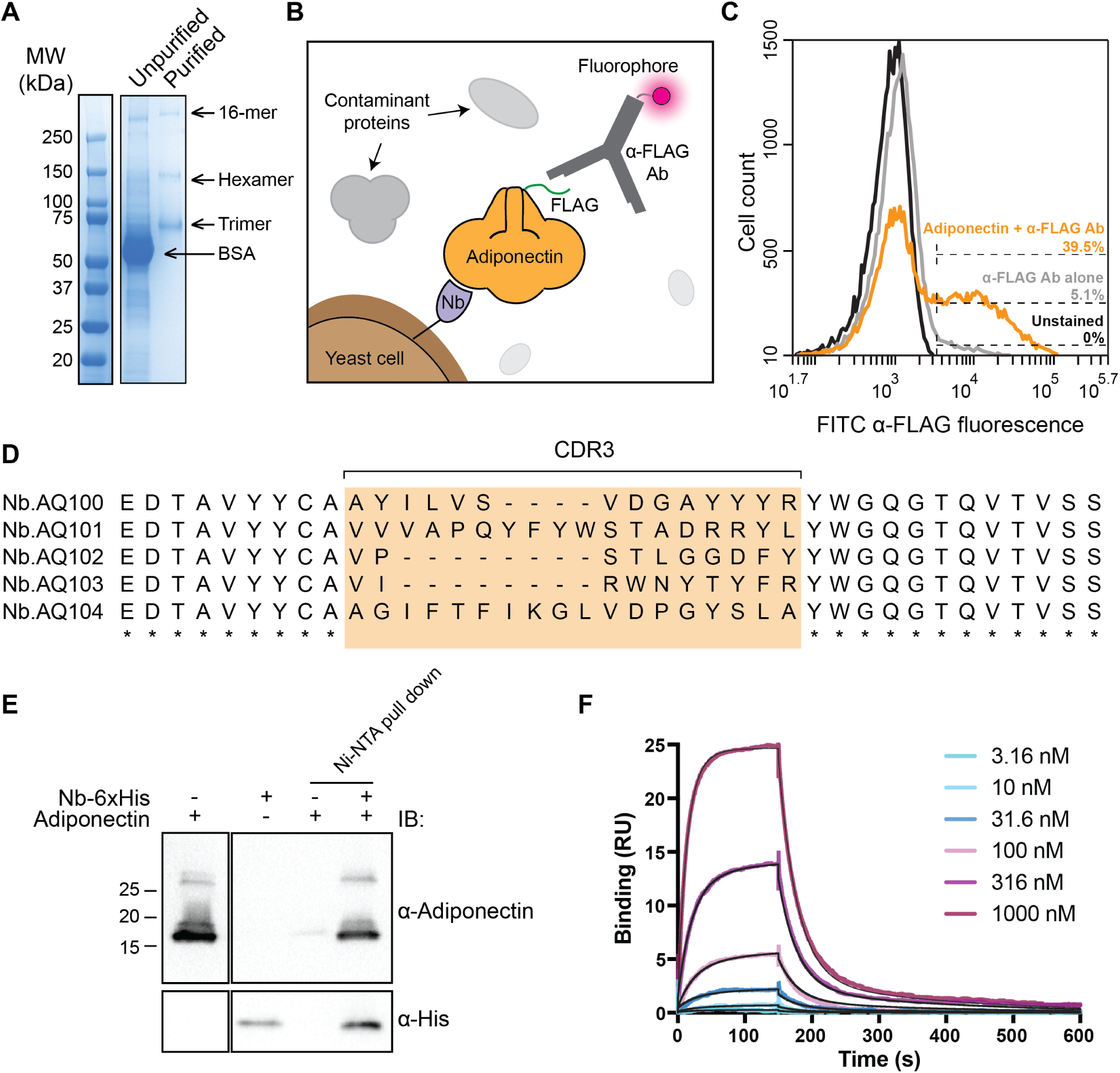
Discovery of nanobodies with non-purified antigen. (A) Conditioned medium containing adiponectin (left lane) was used for selection of nanobodies. It shows a complex mixture of proteins as assessed by SDS-PAGE. For reference, purified adiponectin is shown in the right lane. Adiponectin exists as a mix of 16-mers, hexamers, and trimers. (B) Schematic of selection process. Fluorescent anti-FLAG antibody was used to specifically mark those yeast cells that display adiponectin-binding nanobodies. (C) Flow cytometry analysis of final clone pool, showing that the library was highly enriched in adiponectin-binding clones. (D) Sequences of five selected clones showed highly diverse CDR3 sequence composition and length. (E) Binding assessed using *in vitro* pull-down with purified adiponectin globular domain. (F) Binding to adiponectin was further confirmed *in vitro* using surface plasmon resonance. Kinetic fit is shown for clone Nb.AQ103.

## Flippase-targeted nanobodies

The function of many proteins is driven by changes in their conformation. Hence, correlating specific protein conformations to cellular function remains a key goal of modern structural and cellular biology. Over the past decade, antibodies have provided a remarkably powerful approach to probe the role of protein conformation in function. Such antibodies are not only specific to a given protein, but only bind to that protein when it adopts a specific conformation. These conformationally-selective antibodies have been leveraged to study G protein-coupled receptors^6^, transmembrane transporters^18,19^, dynamic multiprotein complexes^20^, and enzymes^21^. Conformationally-selective antibodies are critically important tools for studies of these proteins and many others, However, obtaining conformationally-selective nanobodies from immunization has proven to be challenging, often requiring extremely stable protein preparations or cross-linking complex assemblies^22^. We hypothesized that our synthetic library approach could provide a simpler and more rapid platform to address this challenge, producing nanobodies suitable for conformational stabilization of integral membrane proteins. As a prototypical test case, we chose to produce inhibitory nanobodies targeting the multidrug oligosaccharidyl-lipid polysaccharide (MOP) family transporter MurJ, a 14-pass transmembrane protein that functions as a flippase for lipid II, the precursor to peptidoglycan^23,24^. Because of its critical biological role, MurJ is essential for cell wall assembly in most bacteria. Central to its function is conformational exchange, binding the lipid II headgroup on the inner leaflet of the bacterial inner membrane and then releasing it on the periplasmic face. Nanobodies that stabilize a specific conformation of the MurJ would be expected to block lipid II export, thereby preventing cell wall assembly and inducing bacterial lysis (Figure 4).

**Figure 4.**
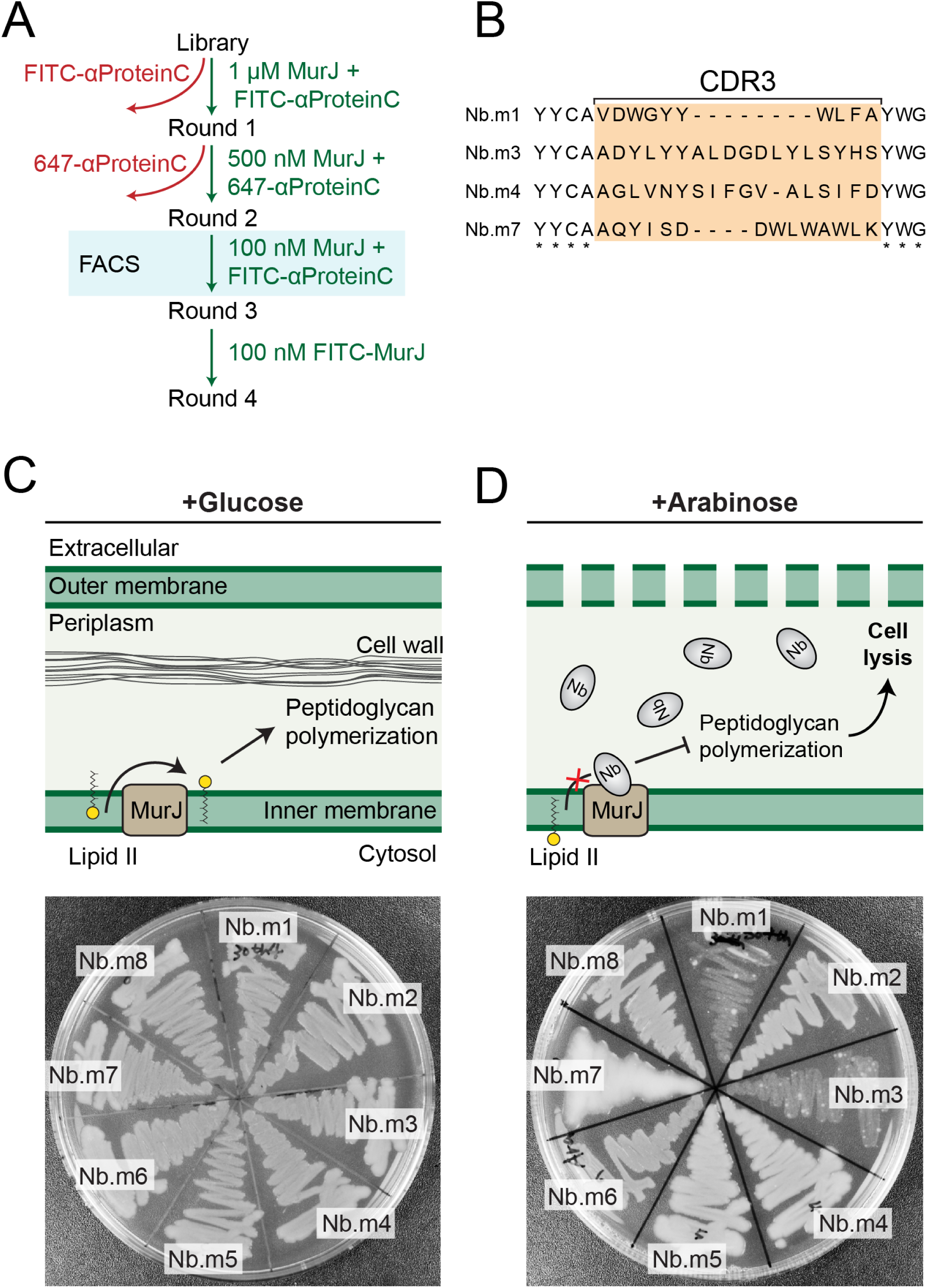
Isolation of flippase-blocking nanobodies. (A) Selection schematic for isolation of MurJ binding nanobodies (B) Sequence analysis of MurJ binding clones, showing divergent CDR3 sequences. (C) Schematic of the role of MurJ as the lipid II flippase in cell wall synthesis under normal conditions. Plating assay with bacteria containing eight nanobodies under non-inducing conditions. (D) Schematic of nanobodies blocking the flippase activity of MurJ leading to cell lysis and bacterial plate with bacteria expressing eight nanobodies against MurJ. Three clones show either complete abrogation of growth (Nb.m1, Nb.m4) or a mucoid phenotype indicating impairment of cell wall assembly (Nb.m7).

To first identify MurJ-targeted nanobodies, we generated and purified *E. coli* MurJ tagged with a carboxyl-terminal Protein C epitope tag (“EDQVDPRLIDGK”). Our yeast library was subjected to two rounds of MACS selection followed by one round FACS using either purified MurJ or fluorescently-labeled anti-Protein C antibody to detect MurJ binding. MurJ-binding yeast were further enriched with an additional round of MACS using fluorescently labeled MurJ (Figure 4A). Because MurJ is essential for cell viability^23^, we predicted that conformationally-selective nanobodies would block transport of lipid II and thereby inhibit bacterial growth. To identify conformationally selective nanobodies targeting the extracellular side of MurJ, we devised a bacterial growth assay in which nanobodies were secreted into the periplasmic space upon induction with arabinose. Tight control of nanobody expression was obtained by glucose-mediated repression of the pBAD promoter. By plating cells on agar containing either arabinose or glucose, the effect of each nanobody clone on bacterial growth could be easily assessed (Figure 4C, D). Out of 115 clones tested, a total of 6 unique nanobodies (5%) completely abrogated cell growth upon induction. This indicates these nanobodies block lipid II flipping by MurJ, likely by preventing conformational exchange or other steps of the transport cycle and suggesting that conformationally-selective membrane-protein targeted nanobodies can be obtained without any animal immunization step.

## Conformationally-selective GPCR-binding nanobodies

While nanobodies have contributed to a broad range of biological fields, they have had a uniquely transformative impact on our understanding of G protein-coupled receptor (GPCR) biology^6^. GPCRs are the largest family of transmembrane receptors in humans, playing essential roles in every aspect of physiology ranging from function of the central nervous system to regulation of metabolism and cardiovascular biology. Unfortunately, GPCRs are among the most challenging proteins to manipulate biochemically. Nanobodies have proven to be invaluable tools in addressing the challenges of GPCR research, enabling profound insights into receptor structure and function. As an example, a llama-derived nanobody, Nb80, that stabilizes the active conformation of the β2-adrenergic receptor (β2AR) enabled determination of the first X-ray crystal structure of the active conformation of a hormone-activated GPCR^4^. The conformational selectivity of Nb80 was also subsequently used to probe the localization of active β2AR in live cells, thereby revealing new paradigms in intracellular GPCR signaling^3^. Another nanobody, Nb35, was instrumental in examining the structure and signaling of the heterotrimeric G protein Gs in complex with the β2AR, enabling determination of the first structure of a GPCR-G protein heterotrimer complex^22^.

Given the broad utility of conformationally-selective antibodies in GPCR research, any *in vitro* platform aiming to surpass the utility of immunization-based methods must be able to generate conformationally selective GPCR binders. Hence, we sought to identify nanobodies that selectively bind the active conformation of β2AR using our synthetic nanobody library platform (Figure 5). Nanobody-displaying yeast were first selected for binding to purified, FLAG-tagged β2AR bound to the agonist BI167107 by MACS. We subsequently introduced a counterselection strategy to deplete undesired clones, which include nanobodies that bind the M1 antibody itself, the Alexa Fluor 647 fluorophore, conformationally invariant epitopes, or the inactive β2AR conformation. For example, in the third MACS round we first depleted clones that bound to β2AR occupied by the high affinity antagonist carazolol prior to enriching clones that bound agonist-occupied receptor. To more rapidly identify the desired agonist-specific nanobodies, we performed two rounds of FACS. Here, the yeast library was incubated simultaneously with both agonist (BI167107) and antagonist (carazolol) occupied β2AR, with each receptor-ligand complex labeled with a specific Alexa fluorophore (Figure 5A – C). Importantly, this procedure where the conformational selectivity of each individual clone can be interrogated directly and selected precisely would be impossible using phage- or mRNA-displayed libraries. Such precise selection using FACS requires a cell-based display system. As an indication that the selected clones bind the desired epitope, we observed that a large fraction were competitive with Nb6B9, a previously affinity matured variant of Nb80 that binds with high affinity to the intracellular side of the β2AR^25^. Finally, we performed a round of MACS with decreased concentration of agonist-bound β2AR to enrich the highest affinity clones (Figure 5C).

**Figure 5.**
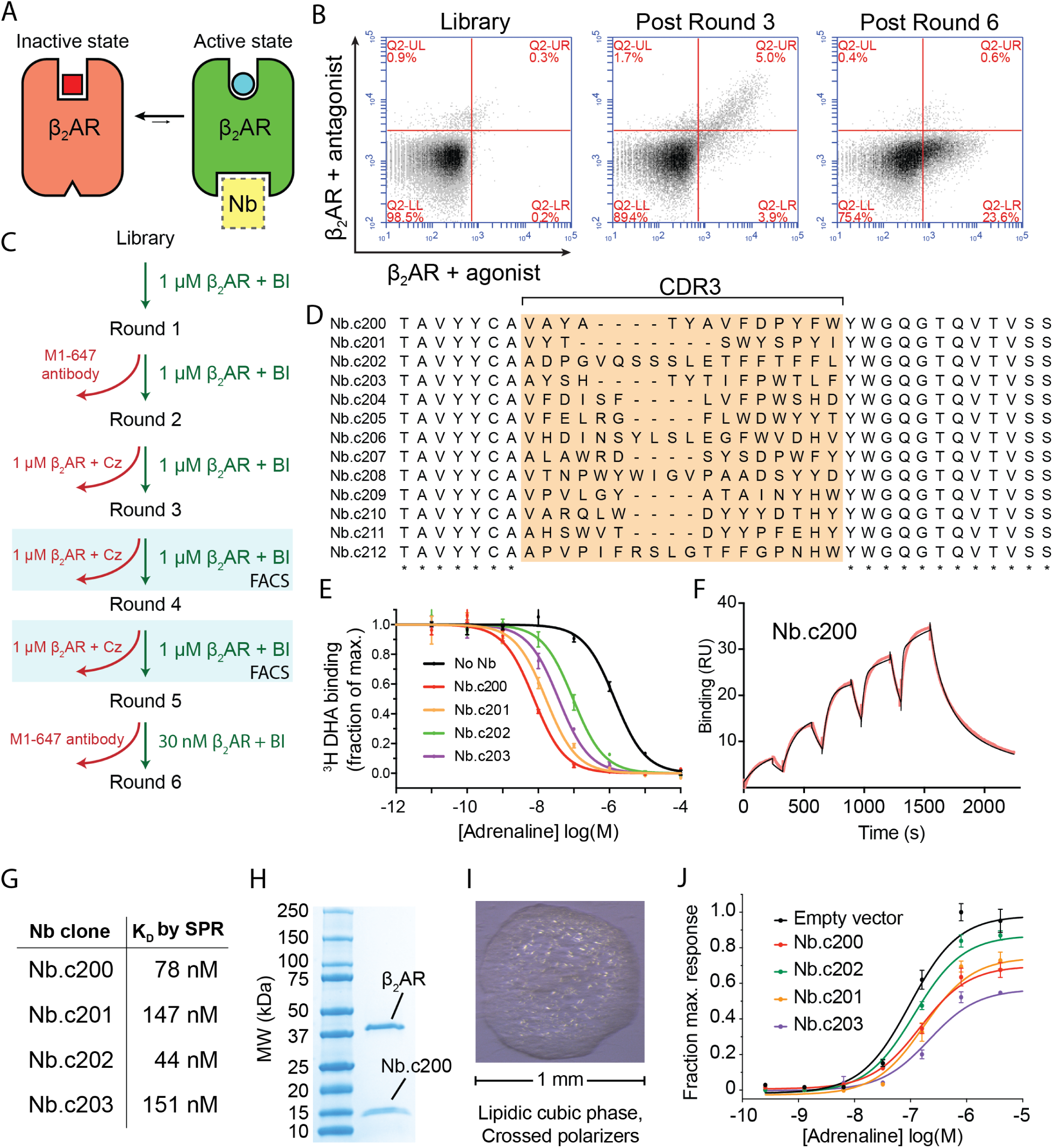
Structural and functional modulator nanobodies targeting a GPCR. (A) β_2_ adrenergic receptor interconverts between ensembles of inactive conformations (red) and active conformations (green). The active state of the receptor can be stabilized by nanobodies (yellow). (B) Results of selection summarized in flow cytometry plots. After FACS selection a significant fraction (23.6%) of clones show agonist-specific binding to the β2AR. (C) Selection schematic for isolation of active-state stabilizing nanobodies. (D) Sequence analysis of β2AR conformationally selective clones, showing highly divergent CDR3 sequences. (E) Radioligand competition binding in nanodiscs confirms that synthetic nanobodies stabilize the active-state conformation β2AR. (F) Single cycle SPR experiment showing measurement of affinity and kinetics for Nb.c200 binding to β2AR-BI167107. Experimental data are in red, and curve fit is in black. (G) Summary of nanobody affinities measured by surface plasmon resonance (see also Figure S5). (H) Nb.c200 and β2AR, co-purified by immobilized metal ion chromatography prior to crystallization. (I) Nb.c200 enabled crystallization of the β2AR using the lipidic cubic phase method. (J) cAMP signaling assay to measure β2AR signaling in the presence or absence of synthetic nanobodies. See also Figure S4.

Sequencing individual clones revealed 13 unique nanobodies that bound agonist-occupied β2AR (Figure 5D**; Figure S4**). We further prioritized clones with on-yeast titrations, which demonstrated that the isolated nanobodies have a range of affinities for the β2AR-BI167107 complex in the low to mid nanomolar range. We anticipate that conformationally-selective nanobodies identified from this library will find utility in biochemical, structural, and cell biological applications. Therefore, we next validated the active-state specific β2AR nanobodies in pharmacological, crystallographic, and cellular signaling experiments. Four clones were selected for further characterization and were expressed and purified from *E. coli*. Each of the four nanobodies increased agonist affinity of β2AR as assessed by a competition radioligand binding assay (Figure 5E). This gold-standard pharmacological assay provides definitive evidence that each nanobody stabilizes the active conformation of β2AR. Next, we determined binding affinities of each nanobody for BI167107-occupied β2AR by surface plasmon resonance (SPR) using single cycle kinetics (Figure 5F, G). Measured affinities ranged from 44 nM to 151 nM, comparing favorably to the most extensively studied llama-derived GPCR-targeting nanobody, Nb80, which binds with roughly 140 nM affinity^25^.

We next tested whether this conformational stabilization is sufficient to enable crystallization of an agonist-bound β2AR. Likely due to conformational dynamics in the agonist-bound state^26^, all previous attempts to crystallize β2AR-BI167107 complex have failed in the absence of an intracellular binding nanobody or the heterotrimeric G protein^4,22,27^. Due to the low throughput of crystallography experiments, we selected only Nb.c200 for crystallization trials. The nanobody readily co-purified with β2AR (Figure 5H), and crystallized easily by the lipidic cubic phase method (Figure 5I). Of note, Nb.c200 enabled crystallization in a range of crystallization conditions rather than just a small subset as is more typical.

Finally, we assessed the potential of our synthetic nanobodies to modulate GPCR signaling in live cells, a key application of llama-derived nanobodies ^3,28^. As with llama-derived nanobodies, we found that our synthetic nanobodies could significantly impair β2AR signaling in response to adrenaline. Among the nanobodies tested, we found that Nb.c203 was the most effective, reducing adrenaline E_max_ by 45%, consistent with the G protein-competitive binding mode predicted from flow cytometry and pharmacological experiments discussed above (Figure 5J).

Taken together, these data show that conformationally-selective nanobodies identified from by our *in vitro* platform recapitulate key features of llama-derived clones. The larger diversity of conformationally-selective clones identified from the synthetic library, however, may enable a larger repertoire of applications; some may perform better as crystallographic chaperones, others may be more stable in the reducing intracellular environment, while others may have increased specificity required for identifying a specific protein in a complex milieu.

## Discussion

Antibodies and their minimal binding fragments have transformed biomedical research. Nanobodies, derived from camelid heavy chain only antibodies, have emerged as powerful alternatives to conventional antibodies. These single domain antibodies have enabled the investigation of protein structure and function and are emerging as potential therapeutic agents. However, despite their broad utility, nanobodies have only been exploited for a subset of potential applications due to challenges associated with nanobody discovery. Foremost among these is the requirement for camel, llama, or alpaca immunization – a slow, expensive process requiring large quantities of purified protein and access to large animal husbandry and veterinary facilities. Moreover, this process frequently yields only small numbers of nanobody clones, many or all of which may not possess the desired activity. Our approach reported here addresses these challenges by providing a fully *in vitro* platform for nanobody discovery based on an engineered library displayed on *S. cerevisiae*.

To assess the utility of our library, we selected nanobodies targeting a diverse array of antigens: soluble proteins, non-purified proteins, and integral membrane proteins of both bacterial and mammalian origin. In each case, our platform enabled identification of nanobody clones capable of binding to the target antigen with the desired activity. To date, all nanobody selection efforts conducted with this library have led to the successful isolation of clones with sub-micromolar affinity, with the exception of an effort targeting FLAG peptide, a short unstructured sequence. Surprisingly, we found that the most effective selections were those with membrane antigens. In the case of the lipid flippase MurJ, several unique transport-blocking clones were obtained, while in the case of the β_2_ adrenergic receptor a total of 12 conformationally-selective clones were isolated. Strikingly, the affinities of our synthetic β2AR binding clones were comparable to or better than the most extensively studied GPCR-targeting nanobody, Nb80, which has an affinity of 140 nM. Moreover, if needed, nanobodies are readily amenable to affinity maturation to improve affinity, even for difficult targets such as GPCRs^25^.

While the approach described here has proven to be a powerful method for developing useful new nanobodies, it is important to acknowledge that like any method it is not without limitation. In particular, we have found that this synthetic library readily yields nanobodies targeting fluorophores and selection reagents such as antibodies or microbeads. While this problem can be avoided, it is critically important to plan selections carefully to avoid enriching such unwanted clones. Frequent alternation of fluorophores and the use of rigorous pre-clearing against selection reagents resolve this issue, but are essential steps that cannot be omitted. A second potential limitation is the fact that this library currently represents only a subset of the types of clones found in the immune repertoire. For instance, the design of this library excludes disulfide-tethered CDR loops, which are found in approximately 25% of llama V_HH_ antibodies and even more frequently in camel-derived nanobodies^29^. It is possible that for some antigens such clones may provide higher affinity binders or offer unique modes of recognition.

Nonetheless, despite these important considerations, our results show that this *in vitro* nanobody discovery platform yields comparable or superior results to animal immunization across a broad range of target classes and intended applications. From soluble protein binders to conformationally-selective GPCR stabilizers, our platform allows straightforward, rapid, and low-cost isolation of nanobodies with desired properties. To facilitate broad deployment of this resource, we have made the library publicly available. Detailed protocols for nanobody binder selection are included in the Supplementary Materials, and can be adapted or modified to enable more diverse applications. We envision that rapid and simple nanobody discovery will be an enabling resource for applications throughout biomedical research.

**Library access:** The synthetic nanobody library is available from the authors for noncommercial use. Please contact Aashish Manglik or Andrew Kruse to request the library.

---

**Figure S1 | Biochemical validation of nanobody clones.** (A – K) Randomly chosen nanobodies were expressed and purified from *E. coli*, then analyzed by size exclusion chromatography to assess monodispersity. (L) SDS-PAGE analysis of nanobody purity following one-step nickel affinity purification.

**Figure S2 | Design of display system.** (A) The display system was engineered using the high affinity SIRPα variant CV1 as a test protein, and its ligand CD47 ectodomain as the staining reagent. A biotin tag is schematized as a glowing red circle. (B) Length of the stalk region determines accessibility of a displayed protein as a function of molecular weight. (C) Analytical flow cytometry plots showing length dependence for two staining reagents: CD47 biotin and α-HA antibody. The 649 amino acid long stalk was used in all nanobody display experiments.

**Figure S3 | Analysis of HSA-targeted nanobodies.** (A) Library design was assessed by monitoring the change in amino acid frequency in CDR3 throughout selection rounds with HSA as the antigen. Few changes were observed, with the only notable trend a modest increase in basic residue frequency and a decline in acidic residue frequency. (B) Assessment of Nb.b201 binding to human serum albumin by surface plasmon resonance, comparison with mouse serum albumin which shows no detectable binding. (C) 2F_o_-Fc composite omit map contoured at 1.5 σ for antigen bound Nb.b201. The structure of both bound (yellow) and free (gray) forms of the nanobody are shown, highlighting structural divergence. (D) 2F_o_-F_c_ composite omit map contoured at 1.5 σ for free Nb.b201.

**Figure S4 | Affinity of β2AR binding nanobodies.** On-yeast titration to estimate affinity of β2AR binding nanobodies. EC_50_ values are summarized in the lower right. Bottom panel shows measurement of conformational selectivity for selected clones as assessed by flow cytometry.

**Table S1 | Sequences of nanobodies.**

**Table S2 | Crystallographic data collection and refinement statistics.**

**Table S3 | Sequences of primers used in this study.**

## Acknowledgments

Financial support for this work was provided by the Vallee Foundation (A.C.K.), the Smith Family Foundation (A.C.K.), and NIH grants 5DP5OD021345 (A.C.K.), 1DP5OD023048 (A.M.), and 1DP5OD023088 (A.M.R.), as well as a supplement to Center for Excellence in Translational Research grant 5U19AI109764 (A.C.K.).

## Methods

### Nanobody library construction

The DNA library of nanobodies was constructed by two-step overlap extension PCR(OE-PCR). A set of ten primers; P1_for, P2_rev, …, P10_rev (Keck Biotechnology Resource Laboratory; **Table S3**), were dissolved at 100 μM concentration and mixed in an equimolar ratio to prepare three mixed pools containing each primer at a concentration of 10 μM. The three mixed pools, ‘mix short’, ‘mix medium’, and ‘mix long’, differed in the P9 primers, using P9a_for, P9b_for, or P9c_for to introduce a CDR3 region of variable length; 7, 11, or 15 randomized residues respectively. 1 μL of each mixed pool at 10 μM concentration and at 5-fold sequential dilutions was subsequently used to prepare 50 μl OE-PCR reactions using Phusion polymerase. 0.4 μM of each primer, or a 25-fold diluted stock, was found to produce optimal yields for OE-PCR for each of the three mixed pools. The full length nanobody DNA product from each pool was mixed in an 1:2:1 molar ratio of short to medium to long CDR3 regions, referred to as the nanobody DNA library pool hereafter, recapitulating the length distribution frequencies observed in camelid V_HH_ domains. Validation of the biochemical tractability of these synthetic nanobodies was carried out in *Escherichia coli* by amplifying the resulting mixture with primers pET26b_NbLib_GA_for and pET26b_NbLib_GA_rev and cloning into pET26b.

The nanobody DNA library pool was successively amplified for yeast transformations with pYDSFor1-pYDSRev1, pyDSFor2-pYDSRev2, and pYDSFor3-pYDSRev2 primers. 500 mL of BJ5465 yeast (MAT**a** *ura352 trp1 leu2Δ1 his3Δ200* pep4::HIS3 *prb1Δ1.6R can1* GAL) were grown to OD_600_ 1.5 and transformed with 245 μg nanobody insert DNA and 50 μg of pYDS649 plasmid, digested with NheI-HF and BamHI-HF (New England BioLabs), using an ECM 830 Electroporator (BTX-Harvard Apparatus). Dilutions of transformed yeast were then plated on –Trp dropout medium as single colonies to obtain an estimate of library diversity.

Three cultures of 5×10^5^ yeast were inoculated in Trp dropout medium (US Biological) and grown overnight at 30 °C for whole library next generation sequencing reactions. From these cultures, PCRs were performed on 8×10^6^ pelleted yeast to amplify nanobody genes using three sets of primers with different barcodes: Batch1For-Batch1Rev, Batch2For-Batch2Rev, and Batch3For- Batch3Rev. A sequencing library was prepared using PrepX (Wafergen) and sequencing was performed by a MiSeq (Illumina) with 10% PhiX. Next generation sequencing of HSA binding nanobody selection rounds was performed similarly starting with 2×10^5^ yeast cells in the initial inoculum and using the primers: Rd1For-Rd1Rev, Rd2For-Rd2Rev, Rd3For-Rd3Rev, Rd4For-Rd4Rev.

Optimization of yeast display plasmid is summarized in **Figure S2**. SIRPα mutant CV1 and CD47 proteins used for the assay were expressed and purified as described previously^30^. Analytical yeast staining experiments were performed as described below, using yeast strain BJ5465 as in other experiments.

### Isolation of nanobody binders from library

#### Human serum albumin

Initially, 2.5 × 10^9^ induced yeast for the first round and 1.4 - 5 × 10^7^ induced yeast for subsequent rounds were washed and resuspended in buffer (20 mM HEPES pH 7.5, 150 mM sodium chloride, 0.1% (w/v) ovalbumin, 1 mM EDTA) and then incubated with anti-Alexa Fluor 647 or anti-FITC microbeads (Miltenyi) at 4 °C for 30 min. Each round of MACS selection began with a pre-clear step which involved passing the yeast, through an LD column (Miltenyi) to remove yeast expressing nanobodies that bound nonspecifically to magnetic beads. After pre-clearing, human serum albumin (HSA) binding nanobodies were enriched over four rounds of MACS selection by staining the yeast alternately with Alexa Fluor 647 or FITC labeled HSA (Sigma) and microbeads, then passing them through an LS column (Miltenyi). During these four selection rounds, yeast were stained with successively lower concentrations of HSA: 1 μM, 250 nM, 75 nM,and 15 nM, in order to enrich for binders with higher affinities. After MACS selection, yeast were plated as single colonies which were picked and grown as clonal populations in a 96 well plate. Following galactose induction of nanobodies, yeast were stained with Alexa Fluor 488 labeled HSA and analyzed by flow cytometry with an Accuri C6 (BD Biosciences) to screen for nanobody binders.

#### Human adiponectin

Human adiponectin bearing N-terminal FLAG tag was expressed in Expi-293 cells (ThermoFisher Scientific) for secretion into the surrounding medium. The approximate concentration of adiponectin in this medium was estimated by dividing the average yield of pure adiponectin obtained for each harvest by volume of supernatant. The adiponectin-containing supernatant taken from the cells was used as the selection reagent, with no actual purification performed.

For the first round of selection, 5 × 10^9^ induced yeast were washed and incubated in buffer (1 × HBS + 2 mM CaCl_2_ + 0.1% BSA + 1.8% maltose) with Alexa647-labelled anti-FLAG M1 and anti-647 microbeads. Clones binding nonspecifically to the staining reagents were removed by passage through an LD column and the remaining yeast from the flow-through were incubated with unpurified adiponectin-FLAG at 500 nM for 30 min at 4 °C. Cells were then stained with anti-FLAG-647 and anti-647 microbeads. Binding clones were enriched via magnetic selection in an LS column, and cultured overnight in –TRP medium at 30 °C. Rounds 2 and 3 of selection were performed similarly with 3 × 10^7^ induced yeast and cells were washed, stained with 647- or FITC-labelled M1, and labelled with anti-Alexa Fluor 647 or anti-FITC microbeads before magnetic separation during rounds 2 and 3, respectively.

Following round 3, induced cells incubated with ~500 nM unpurified adiponectin and stained simultaneously with M1-647 and M1-FITC to differentiate dye-, antibody-, and adiponectin-binding clones. FACS was used to enrich for cells that exhibited adiponectin-dependent 647- and FITC staining. Nanobodies from individual clones were sequenced and purified and adiponectin binding was confirmed and characterized by SPR with purified adiponectin.

#### Bacterial lipid II flippase MurJ

To isolate MurJ^A29C^ binding nanobodies, two rounds of MACS were performed with purified MurJ, tagged with a carboxyl-terminal Protein C sequence, and FITC or Alexa Fluor 647 labeled anti-Protein C antibody in selection buffer containing 20 mM HEPES pH 7.5, 250 mM NaCl, 0.01% lauryl maltose neopentyl glycol (L-MNG, Anatrace), 0.001% cholesterol hemisuccinate (CHS, Anatrace), 2 mM CaCl_2_, and 5 mM maltose. Prior to each round of MACS, the yeast library was precleared, as detailed above, to remove binders to fluorophore labeled anti-Protein C antibody or the magnetic beads. Precleared yeast were incubated with 1 μM or 500 nM of MurJ for selection rounds 1 and 2, respectively, along with anti-Protein C antibody and anti-Alexa Fluor 647 or anti-FITC microbeads and applied onto LS column. For the third round of selection, yeast were incubated with 100 nM MurJ, FITC labeled anti-Protein C and Alexa Fluor 647 labeled anti-HA antibody to select for clones that demonstrated staining consistent with both binding and expression, respectively, using FACS. For the fourth round MACS selection, 100 nM MurJ, directly labeled with FITC, was used as a final enrichment step. Nanobodies from the 3^rd^ and 4^th^ rounds of selection were cloned into the bacterial expression vector PMT167 for periplasmic expression. In both vectors nanobodies were under the control of a pBAD promoter which is activated by arabinose and repressed by glucose. The resulting nanobody library was electroporated into TB28 *E. coli*. Individual clones were streaked onto LB plates supplemented with 25 μg/mL chloramphenicol and 0.2% arabinose or 0.2% glucose.

#### Human β_2_-adrenergic receptor

To isolate agonist-specific β_2_AR binding nanobodies, two rounds of MACS were performed with 1 μM purified, FLAG tagged β_2_AR bound to the high affinity agonist BI167107 and FITC or Alexa Fluor 647 labeled anti-FLAG antibody in a selection buffer containing 20 mM HEPES pH 7.5, 100 mM NaCl, 0.1% dodecylmaltoside (DDM, Anatrace), 0.01% CHS, 5 mM CaCl_2_, and 10 mM maltose. For round 2 of selection the yeast library was precleared, as detailed above, with anti-FLAG antibody labeled with Alexa Fluor 647 to deplete antibody or fluorophore binders. Round 3 utilized MACS again, but counterselection was performed against β_2_AR bound to the high affinity antagonist carazolol to deplete nanobody clones that bind conformationally invariant or inactive-state epitopes. In subsequent rounds, we utilized FACS to more specifically select nanobodies that selectively bind β_2_AR in the active conformation. In round 4, yeast were simultaneously incubated with 1 μM β_2_AR-BI167107 complex labeled with Alexa Fluor 647 NHS ester and 1 μM β_2_AR-carazolol complex labeled with Alexa Fluor 488 NHS ester. Yeast displaying selectivity for agonist (high Alexa Fluor 647 signal and low Alexa Fluor 488 signal) were collected, expanded in growth media, induced and subjected to another round of FACS. The conformational selection procedure was repeated as in round 4, however the fluorophores coupled to agonist and antagonist were switched to deplete non-specific clones. A final MACS round was performed at 30 nM β_2_AR-BI167107 to isolate the highest affinity nanobodies. Nanobodies from individual clones were subsequently sequenced and purified for biochemical and biophysical studies.

### Protein expression and purification

Adiponectin gene was cloned into pTARGET vector with an HA signal peptide that directs secretion into the medium and a FLAG tag in front of the main gene, so the mature protein has an N-terminal FLAG epitope with Asp at the very first residue. A 293-Expi stable cell line was created to express FLAG-Adiponectin in the medium and was maintained into a bioreactor (CELLine Flasks, Wheaton). The cell medium of the lower chamber is composed of Expi293 Expression Medium (Thermo Fisher) implemented with 10% FBS (Zen-Bio), 10 μg/mL gentamicin (VWR). The medium of the upper chamber is FreeStyle 293 Expression Medium (Life Technologies), supplemented with 10 ug/mL gentamicin (VWR). Cells from the lower chamber were harvested by centrifugation at 4000 g for 10 minutes. The supernatant was diluted 1:1 with HBS and CaCl_2_ 2 mM was added before loading on to a FLAG resin. FLAG resin was first washed with 300 mM NaCl, 20 mM Hepes pH 7.5, CaCl_2_ 2 mM and then with 150 mM NaCl, 20 mM Hepes pH 7.5, CaCl_2_ 2 mM; elution was performed with 150 mM NaCl, 20 mM Hepes pH 7.5, FLAG peptide 0.2 mg/mL, EDTA 5 mM. Adiponectin was dialyzed in HBS overnight.

Human β_2_AR fused to an amino-terminal hemagglutinin signal peptide and FLAG-tag as well as a carboxy-terminal 1D4 tag was purified from infected *Sf9* cells grown in ESF 921 media (Expression Systems). Cells were collected 64 hours after baculoviral infection and stored at −80 °C. For receptor preparations used during nanobody selections, purification was performed using ligand-affinity chromatography as described previously^31^. For biophysical assays, we used a simpler purification method omitting ligand-affinity chromatography. Cells were lysed in a buffer comprised of 20 mM HEPES pH 7.5, 2 mM MgCl_2_, 2 μl benzonase, and 1 μM ICI-118,551. The resulting lysate was centrifuged, and the pellets were resuspended in solubilization buffer containing 20 mM HEPES pH7.5, 200 mM NaCl, 10% glycerol, 1% - MNG, 0.1% CHS, and 1 μM ICI-118,551 followed by incubation at 4 °C for 2 h. The insoluble fraction was separated by centrifugation at 39,000x g for 20 min, and the supernatant was loaded over 5 mL of homemade M1-FLAG resin. FLAG resin was subsequently washed with 20 mM HEPES pH7.5, 200 mM NaCl, 10% glycerol, 0.1% MNG, and 0.01% CHS. β_2_AR was eluted in the same buffer with a lower concentration of detergent and CHS (0.01% and 0.001%, respectively) supplemented with 5 mM EDTA and 0.2 mg/mL FLAG peptide. Receptor was further purified by size exclusion chromatography in the presence of agonist BI167107. For crystallographic experiments, β_2_AR fused to an amino-terminal T4 lysozyme^32^ was similarly purified.

Isolated nanobody genes were cloned into the periplasmic expression vector pET26b containing a C-terminal 6×histidine tag and expressed from BL21 (DE3) *E. coli*. For each purification, *E. coli* were grown in Terrific Broth (Research Products International) and induced after reaching OD_600_ 0.6-0.8 with 1 mM IPTG (Gold Biotechnology) at 25 °C for about 15 hours. Induced cells were pelleted and resuspended in 100 mL of buffer consisting of 0.5 M sucrose, 0.2 M Tris pH 8, 0.5 mM EDTA and osmotically shocked by adding 200 mL of water with 45 minutes of stirring, releasing periplasmic nanobody. The lysate was brought to a concentration of 150 mM NaCl, 2 mM MgCl_2_, and 20 mM imidazole and then centrifuged 20,000 × G for 20 min at 4 °C. The supernatant was applied to a gravity column with 3 mL of Ni Sepharose 6 Fast Flow (GE Healthcare). The resin was washed with a high salt buffer (20 mM HEPES pH 7.5, 500 mM NaCl, 20 mM Imidazole) and then washed with NH7.5 buffer (20 mM HEPES pH7.5, 100 mM NaCl) + 20 mM imidazole and nanobodies were eluted with NH7.5 + 400 mM imidazole. Nanobodies used for crystallography were treated to carboxypeptidase A (Sigma) and B (Roche) to remove the histidine tag, dialyzed into HBS buffer (10 mM HEPES pH 7.4, 150 mM NaCl), and co-purified with their respective binding partners over an S200 size-exclusion column (GE Healthcare).

### Characterization of nanobody binding

Binding of Nb.b201 with HSA or MSA (Sigma) was qualitatively assessed by mixing the proteins in a 1.2:1 ratio, respectively, and separating through a Superdex 200 10/300 gel filtration column (GE Healthcare) in HBS buffer.

For SPR of Nb.b201, EZ-Link NHS-PEG4-Biotin (Thermo Fisher Scientific) labeled recombinant HSA from *S. cerevisiae* (HSAr; Sigma) or MSA were immobilized to a Series S Sensor Chip SA using a Biacore T200 (GE Healthcare). Dilutions of Nb.b201 in running buffer (10 mM HEPES pH 7.5, 150 mM NaCl, 0.03% Tween) were made and the sample was injected at a flow rate of 30 μl/minute with a contact time of 120 s and dissociation time of 300 s. For SPR of β2AR binding nanobodies, β2AR was labeled with NHS-PEG4-Biotin and immobilized on a Sensor Chip CAP (GE Healthcare). Dilutions of β2AR were made in running buffer (10 mM HEPES pH 7.5, 150 mM NaCl, 0.01% MNG, 0.001% CHS, 20 nM BI167107) and the sample was injected at a flow rate of 30 μl/min with a contact time of 240 s and dissociation time of 700 s.

For SPR of Nb.AQ103, EZ-Link NHS-PEG4-Biotin (Thermo Fisher Scientific) labeled FLAG-Adiponectin was immobilized to a Series S Sensor Chip SA using a Biacore T200 (GE Healthcare). Dilutions of Nb.AQ103 in running buffer (20 mM HEPES pH 7.5, 150 mM NaCl, 0.05% Tween) were made and the sample was injected at a flow rate of 30 μl/minute with a contact time of 150 s and dissociation time of 500 s.

For SEC-MALS, a 1.4:1 molar mix of Nb.b201 and HSAr, as well as Nb.b201 and HSAr alone controls, were injected into an AdvanceBio 300 Å column (Agilent) using an Agilent 1260 Infinity Isocratic Liquid Chromatography System and characterized using a Wyatt DAWN HELEOS II Multi-Angle static Light Scattering detector.

For HIS-pull down assay, 5 μM Nb.AQ103 was incubated with Ni^2+^ resin at 4 °C for 1 h, then the excess was removed with wash with HBS and then incubated with adiponectin globular domain at 4 °C for 1 h. The reaction was washed with HBS and imidazole 20 mM and eluted with HBS in presence of 250 mM imidazole.

### Validation of conformationally-selective β_2_AR nanobodies

For radioligand binding studies, β2AR was reconstituted into high-density lipoprotein (HDL) nanodiscs containing a 3:2 molar ratio of POPC/POPG as described previously^33^. Binding experiments were performed with 150 fmol β2AR. Saturation-binding was first performed using ^3^H-dihydroalprenolol (^3^H-DHA; PerkinElmer), yielding a K_D_ of 0.7 nM. Competition binding experiments to measure adrenaline affinity in the presence and absence of nanobodies were carried out in a binding buffer comprised of 20 mM HEPES pH 7.5, 150 mM NaCl, 0.1% BSA, and 1 mM EDTA containing 2 nM ^3^H-DHA, 150 fmol β_2_AR, and a range of adrenaline concentrations 10^-4^ - 10^-11^ M. Nanobodies were used at a final concentration of 5 μM. Reactions were incubated for 1.5 hr at room temperature prior to rapid filtration with a Brandel 96-well harverster onto a filter pre-treated with 0.1% polyethyleneimine. Bound ^3^H-DHA was measured by liquid scintillation counting. Measurements were performed in triplicate and are expressed as mean +/- SEM.

### β2AR cAMP signaling assay

β2AR signaling was measured using a transcriptional CRE-SEAP (secreted embryonic alkaline phosphatase) reporter to indirectly measure cAMP production^34^. Briefly, HEK293T cells were seeded at 3.3×10^4^ cells/well in 96-well plates the day before transfection in 200 μL/well of DMEM (+4.5 g/L glucose, +L-glutamine, no sodium pyruvate; Life Technologies) supplemented with 10% FBS. The following day, medium was aspirated from the cells and replaced with 50 μL of serum-free DMEM, and cells at 70% confluency were transfected in triplicate with Lipofectamine 2000 (Thermo Fisher Scientific) according to the manufacturer’s instructions. Each well was transfected with 10 ng of pcDNA3.1(+) encoding β2AR, 20 ng of CRE-SEAP reporter plasmid (BD Biosciences), and 20 ng of pEGFP-N1. Plasmids included conformationally-selective β2AR nanobodies (Nb.c200 - c203), a negative control nanobody (Nb.BV025), and the pEGFP-N1 empty vector. After six hours of incubation at 37 °C, the transfection mix was aspirated from the cells and 200 μL of serum-free DMEM was added with the indicated final concentrations of adrenaline. The cells were incubated at 37 °C for 48 hours, then at 70 °C for 2 hours. To determine SEAP activity, the substrate 4-methylumbelliferyl phosphate (Sigma Aldrich) was prepared at 1.2 mM in 2 M diethanolamine bicarbonate pH 10 and mixed with an equal volume of cell supernatant at room temperature for 10 minutes. Fluorescence was measured on an EnVision 2103 Multilabel Reader (Perkin Elmer) with an excitation wavelength of 360 nm and an emission wavelength of 449 nm. For each condition, the baseline fluorescence value was determined from cells not treated with adrenaline and was subtracted from every value in the data set. β2AR signaling was calculated as a fraction of the maximum observed response (pEGFP-N1 empty vector) and plotted using GraphPad Prism.

### Protein crystallization

Nb.b201 for crystallographic study was purified from *E. coli* as described above. The Nb.b201 complex with HSA was prepared by mixing human serum albumin (Sigma) with Nb.b201 at a ratio of 1:1.15, followed by size exclusion chromatography purification. Purified complex was concentrated to 120 mg/mL and mixed in 200 nL + 200 nL drops with Morpheus HT-96 screen (Molecular Dimensions). Crystals were grown in a sitting drop format, and were obtained directly from the screen without further optimization. The precipitant solution consisted of 0.02 M each of 1,6-hexanediol, 1-butanol, 1,2-propanediol, 2-propanol, 1,4-butanediol, 1,3-propanediol; 100 mM Tris(base)/BICINE pH 8.5, 20% (v/v) PEG 500 MME, 10% w/v PEG 20,000. Crystals of Nb.b201 alone were obtained when the complex was mixed in a 1:1 drop with a solution comprised of 4.0 M Potassium formate, 0.1 M BIS-TRIS propane pH 9.0, 2% w/v Polyethylene glycol monomethyl ether 2,000. Crystals were soaked with 20% glycerol as a cryoprotectant prior to flash freezing in liquid nitrogen.

For β2AR/Nb.c200 crystallization, the two-syringe mixing method was used to reconstitute the sample in lipidic cubic phase^35^. Crystals were obtained from MemMeso LCP screen (Molecular Dimensions) conditions consisting of 100 mM MES pH 6, 100 mM NaCl, 100 mM MgCl_2_, and 40% PEG 200. Crystals were also obtained in a variety of additional conditions.

### Data collection and structure refinement

Data collection was performed at Advanced Photon Source GM/CA beamlines 23ID-B and 23ID-D. Diffraction data were collected at an energy of 12 keV with 0.2 sec exposure per frame. Each frame covered a 0.2 degree oscillation, and beam intensity was attenuated 100 to 1000 fold depending on crystal size. The structure of Nb.b201 in isolation was solved by molecular replacement in *Phaser* using the structure of the β_2_ adrenergic receptor binding nanobody Nb80 as a search model (PDB ID: 3P0G)^4^. The structure of the HSA:Nb.b201 (PDB: 5VNW) complex was solved by molecular replacement using the structure of Nb.b201 (PDB: 5VNV) and the structure of HSA (PDB ID: 3JRY)^36^ as search models.

Structural refinement for Nb.b201 in isolation was performed by alternate manual building in Coot^37^ and reciprocal space refinement in *Phenix* ^38^. In the final stages, TLS refinement was used to model anisotropic B factors. Refinement for HSA:Nb.b201 complex was performed similarly, with the additional inclusion of non-crystallographic symmetry restraints in the first two rounds of refinement. As in the case of Nb.b201 in isolation, TLS refinement was used in final stages of refinement. Crystallographic statistics are summarized in **Table S2**. Crystallographic data analysis was performed with *xds* and *phenix.refine*, using standard metrics to assess structure quality. Full details of crystallographic statistics are summarized in **Table S2**.

## References

1 Hamers-Casterman, C. et al. Naturally occurring antibodies devoid of light chains. Nature 363, 446–448, doi:10.1038/363446a0 (1993).

2 Muyldermans, S. Nanobodies: natural single-domain antibodies. Annu Rev Biochem 82, 775–797, doi:10.1146/annurev-biochem-063011-092449 (2013).

3 Irannejad, R. et al. Conformational biosensors reveal GPCR signalling from endosomes. Nature 495, 534–538, doi:10.1038/nature12000 (2013).

4 Rasmussen, S. G. et al. Structure of a nanobody-stabilized active state of the beta(2) adrenoceptor. Nature 469, 175–180, doi:10.1038/nature09648 (2011).

5 Staus, D. P. et al. Allosteric nanobodies reveal the dynamic range and diverse mechanisms of G-protein-coupled receptor activation. Nature 535, 448–452, doi:10.1038/nature18636 (2016).

6 Manglik, A., Kobilka, B. K. & Steyaert, J. Nanobodies to Study G Protein-Coupled Receptor Structure and Function. Annu Rev Pharmacol Toxicol 57, 19–37, doi:10.1146/annurev-pharmtox-010716-104710 (2017).

7 Moutel, S. et al. NaLi-H1: A universal synthetic library of humanized nanobodies providing highly functional antibodies and intrabodies. Elife 5, doi:10.7554/eLife.16228 (2016).

8 Adams, J. J. & Sidhu, S. S. Synthetic antibody technologies. Curr Opin Struct Biol 24, 1–9, doi:10.1016/j.sbi.2013.11.003 (2014).

9 Kayushin, A., Korosteleva, M. & Miroshnikov, A. Large-scale solid-phase preparation of 3′- unprotected trinucleotide phosphotriesters--precursors for synthesis of trinucleotide phosphoramidites. Nucleosides Nucleotides Nucleic Acids 19, 1967–1976, doi: 10.1080/15257770008045471 (2000).

10 Kayushin, A. L. et al. A convenient approach to the synthesis of trinucleotide phosphoramidites--synthons for the generation of oligonucleotide/peptide libraries. Nucleic Acids Res 24, 3748–3755 (1996).

11 Boder, E. T. & Wittrup, K. D. Yeast surface display for screening combinatorial polypeptide libraries. Nat Biotechnol 15, 553–557, doi:10.1038/nbt0697-553 (1997).

12 Rakestraw, J. A., Sazinsky, S. L., Piatesi, A., Antipov, E. & Wittrup, K. D. Directed evolution of a secretory leader for the improved expression of heterologous proteins and full-length antibodies in Saccharomyces cerevisiae. Biotechnol Bioeng 103, 1192–1201, doi:10.1002/bit.22338 (2009).

13 Orlean, P. Architecture and biosynthesis of the Saccharomyces cerevisiae cell wall. Genetics 192, 775–818, doi:10.1534/genetics.112.144485 (2012).

14 Makrides, S. C. et al. Extended in vivo half-life of human soluble complement receptor type 1 fused to a serum albumin-binding receptor. J Pharmacol Exp Ther 277, 534–542 (1996).

15 Van Roy, M. et al. The preclinical pharmacology of the high affinity anti-IL-6R Nanobody(R) ALX-0061 supports its clinical development in rheumatoid arthritis. Arthritis Res Ther 17, 135, doi:10.1186/s13075-015-0651-0 (2015).

16 Tijink, B. M. et al. Improved tumor targeting of anti-epidermal growth factor receptor Nanobodies through albumin binding: taking advantage of modular Nanobody technology. Mol Cancer Ther 7, 2288–2297, doi:10.1158/1535-7163.MCT-07-2384 (2008).

17 Kim, C. C., Wilson, E. B. & DeRisi, J. L. Improved methods for magnetic purification of malaria parasites and haemozoin. Malar J 9, 17, doi:10.1186/1475-2875-9-17 (2010).

18 Jiang, X. et al. Crystal structure of a LacY-nanobody complex in a periplasmic-open conformation. Proc Natl Acad Sci U S A 113, 12420–12425, doi:10.1073/pnas.1615414113 (2016).

19 Smirnova, I. et al. Transient conformers of LacY are trapped by nanobodies. Proc Natl Acad Sci U S A 112, 13839–13844, doi:10.1073/pnas.1519485112 (2015).

20 Li, L. et al. Crystal structure of a substrate-engaged SecY protein-translocation channel. Nature 531, 395–399, doi:10.1038/nature17163 (2016).

21 Wu, S. et al. Fabs enable single particle cryoEM studies of small proteins. Structure 20, 582–592, doi:10.1016/j.str.2012.02.017 (2012).

22 Rasmussen, S. G. et al. Crystal structure of the beta2 adrenergic receptor-Gs protein complex. Nature 477, 549–555, doi:10.1038/nature10361 (2011).

23 Sham, L. T. et al. Bacterial cell wall. MurJ is the flippase of lipid-linked precursors for peptidoglycan biogenesis. Science 345, 220–222, doi:10.1126/science.1254522 (2014).

24 Kuk, A. C., Mashalidis, E. H. & Lee, S. Y. Crystal structure of the MOP flippase MurJ in an inward-facing conformation. Nat Struct Mol Biol 24, 171–176, doi:10.1038/nsmb.3346 (2017).

25 Ring, A. M. et al. Adrenaline-activated structure of beta2-adrenoceptor stabilized by an engineered nanobody. Nature 502, 575–579, doi:10.1038/nature12572 (2013).

26 Manglik, A. & Kobilka, B. The role of protein dynamics in GPCR function: insights from the beta2AR and rhodopsin. Curr Opin Cell Biol 27, 136–143, doi:10.1016/j.ceb.2014.01.008 (2014).

27 Rosenbaum, D. M. et al. Structure and function of an irreversible agonist-beta(2) adrenoceptor complex. Nature 469, 236–240, doi:10.1038/nature09665 (2011).

28 Staus, D. P. et al. Regulation of beta2-adrenergic receptor function by conformationally selective single-domain intrabodies. Mol Pharmacol 85, 472–481, doi:10.1124/mol.113.089516 (2014).

29 Vu, K. B., Ghahroudi, M. A., Wyns, L. & Muyldermans, S. Comparison of llama VH sequences from conventional and heavy chain antibodies. Mol Immunol 34, 1121–1131 (1997).

30 Weiskopf, K. et al. Engineered SIRPalpha variants as immunotherapeutic adjuvants to anticancer antibodies. Science 341, 88–91, doi:10.1126/science.1238856 (2013).

31 Manglik, A. et al. Structural Insights into the Dynamic Process of beta2-Adrenergic Receptor Signaling. Cell 161, 1101–1111, doi:10.1016/j.cell.2015.04.043 (2015).

32 Zou, Y., Weis, W. I. & Kobilka, B. K. N-terminal T4 lysozyme fusion facilitates crystallization of a G protein coupled receptor. PLoS One 7, e46039, doi:10.1371/journal.pone.0046039 (2012).

33 Whorton, M. R. et al. A monomeric G protein-coupled receptor isolated in a high-density lipoprotein particle efficiently activates its G protein. Proc Natl Acad Sci U S A 104, 7682–7687, doi:10.1073/pnas.0611448104 (2007).

34 Liberles, S. D. & Buck, L. B. A second class of chemosensory receptors in the olfactory epithelium. Nature 442, 645–650, doi:10.1038/nature05066 (2006).

35 Caffrey, M. & Cherezov, V. Crystallizing membrane proteins using lipidic mesophases. Nat. Protocols 4, 706–731 (2009).

36 Hein, K. L. et al. Crystallographic analysis reveals a unique lidocaine binding site on human serum albumin. J Struct Biol 171, 353–360, doi:10.1016/j.jsb.2010.03.014 (2010).

37 Emsley, P. & Cowtan, K. Coot: model-building tools for molecular graphics. Acta Crystallogr D Biol Crystallogr 60, 2126–2132, doi:10.1107/s0907444904019158 (2004).

38 Adams, P. D. et al. PHENIX: a comprehensive Python-based system for macromolecular structure solution. Acta Crystallogr D Biol Crystallogr 66, 213–221, doi:10.1107/S0907444909052925 (2010).

